# Modelling the impact of the macroalgae *Asparagopsis taxiformis* on rumen microbial fermentation and methane production

**DOI:** 10.1101/2020.11.09.374330

**Authors:** Rafael Muñoz-Tamayo, Juana C. Chagas, Mohammad Ramin, Sophie J. Krizsan

## Abstract

**Background:** The red macroalgae Asparagopsis taxiformis is a potent natural supplement for reducing methane production from cattle. A. taxiformis contains several anti-methanogenic compounds including bromoform that inhibits directly methanogenesis. The positive and adverse effects of A. taxiformis on the rumen microbiota are dose-dependent and operate in a dynamic fashion. It is therefore key to characterize the dynamic response of the rumen microbial fermentation for identifying optimal conditions on the use of A. taxiformis as a dietary supplement for methane mitigation. Accordingly, the objective of this work was to model the effect of A. taxiformis supplementation on the rumen microbial fermentation under in vitro conditions. We adapted a published mathematical model of rumen microbial fermentation to account for A. taxiformis supplementation. We modelled the impact of A. taxiformis on the fermentation and methane production by two mechanisms, namely (i) direct inhibition of the growth rate of methanogens by bromoform and (ii) hydrogen control on sugars utilization and on the flux allocation towards volatile fatty acids production. We calibrated our model using a multi-experiment estimation approach that integrated experimental data with six macroalgae supplementation levels from a published in vitro study assessing the dose-response impact of A. taxiformis on rumen fermentation.

**Results:** our model captured satisfactorily the effect of A. taxiformis on the dynamic profile of rumen microbial fermentation for the six supplementation levels of A. taxiformis with an average determination coefficient of 0.88 and an average coefficient of variation of the root mean squared error of 15.2% for acetate, butyrate, propionate, ammonia and methane.

**Conclusions:** our results indicated the potential of our model as prediction tool for assessing the impact of additives such as seaweeds on the rumen microbial fermentation and methane production in vitro. Additional dynamic data on hydrogen and bromoform are required to validate our model structure and look for model structure improvements. We expect this model development can be useful to help the design of sustainable nutritional strategies promoting healthy rumen function and low environmental footprint.

## 1. Background

Some macroalgae (seaweeds) have the potential to be used as natural supplement for reducing methane (CH_4_) production from cattle (Wang *et al*., 2008; Dubois *et al*., 2013; Maia *et al*., 2016). This anti-methanogenic activity adds value to the nutritional and healthy promoting properties of macroalgae in livestock diets (Evans and Critchley, 2014; Makkar *et al*., 2016). The species of the red macroalgae *Asparagopsis* have proven a strong anti-methanogenic effect both *in vitro* (Machado *et al*., 2014) and *in vivo* (Roque *et al*., 2019). In particular, *Asparagopsis taxiformis* appears as the most potent species for methane mitigation with studies reporting a reduction in enteric methane up to 80% in sheep (Li *et al*., 2016) and up to 80% and 98% in beef cattle (Kinley *et al*., 2020; Roque *et al*., 2020). The anti-methanogenic power of *A. taxiformis* results from the action of its multiple secondary metabolites with antimicrobial activities, being bromoform the most abundant anti-methanogenic compound (Machado *et al*., 2016b). It should be said, however, that despite the promising anti-methanogenic capacity of bromoform, the feasibility of supplying bromoform-containing macroalgae requires a global assessment to insure safety of feeding and low environmental footprint from the algae processing, since bromoform can be toxic to the environment and can impair human health (Beauchemin *et al*., 2020).

Bromoform is released from specialised gland cells of the macroalage (Paul *et al*., 2006) in to the culture medium. The mode of action of the anti-methanogenic activity of bromoform is similar to that described for bromochloromethane (Denman *et al*., 2007), following the mechanism suggested for halogenated hydrocarbons (Wood *et al*., 1968; Czerkawski and Breckenridge, 1975). Accordingly, bromoform inhibits the cobamid dependent methyl-transfer reactions that lead to methane formation. In addition to the direct effect on the methanogenesis, the antimicrobial activity of *A. taxiformis* impacts the fermentation profile *(e.g*., acetate:propionate ratio) and the structure of the rumen microbiota *(e.g*., the relative abundance of methanogens) (Machado *et al*., 2018; Roque *et al*., 2019). Fermentation changes may have detrimental effects on animal health and productivity (Chalupa, 1977; Li *et al*., 2016). Detrimental effects might include deterioration of the ruminal mucosa and the transfer of bromoform to tissues, blood and milk. Previous studies have not detected bromoform in animal tissues (Li *et al*., 2016; Kinley *et al*., 2020; Roque *et al*., 2020). The positive and adverse effects of *A. taxiformis* on the rumen microbiota are dose-dependent (Machado *et al*., 2016a) and operate in a dynamic fashion. It is therefore key to characterize the dynamic response of the rumen microbial fermentation for identifying optimal conditions on the use of the *A. taxiformis* as a dietary supplement for methane mitigation. The development of dynamic mathematical models provides valuable tools for the assessment of feeding and mitigation strategies (Ellis *et al*., 2012) including developments in the manipulation of the flows of hydrogen to control rumen fermentation (Ungerfeld, 2020). Progress on rumen modelling including a better representation of the rumen microbiota and the representation of additives on the fermentation is central for the deployment of predictive tools that can guide microbial manipulation strategies for sustainable livestock production (Huws *et al*., 2018). Accordingly, the objective of this work was to model the effect of *A. taxiformis* supplementation on the dynamics of rumen microbial fermentation under *in vitro* conditions. We adapted a published rumen fermentation model (Muñoz-Tamayo *et al*., 2016) to account for the impact of *A. taxiformis* on rumen fermentation and methane production evaluated *in vitro* at six supplementation levels (Chagas *et al*., 2019).

## 2. Methods

### 2.1. Experimental data

Model calibration was performed using experimental data from an *in vitro* batch study assessing the dose-response impact of *A. taxiformis* on fermentation and methane production (Chagas *et al*., 2019). In such a study, *A. taxiformis* with 6.84 mg/g DM bromoform concentration was supplemented at six treatment levels (0, 0.06, 0.13, 0.25, 0.5, and 1.0 % of diet organic matter; OM). All experimental treatments were composed of a control diet consisted of timothy grass (*Phleum pratense*), rolled barley (*Hordeum vulgare*), and rapeseed (*Brassica napus*) meal in a ratio of 545:363:92 g/kg diet dry matter (DM) presenting chemical composition as 944 g/kg OM, 160 g/kg crude protein (CP) and 387 g/kg neutral detergent fiber (NDF). Prior to each *in vitro* incubation, dried individual ingredients milled at 1 mm were weighted into serum bottles totalizing 1000 mg substrate on DM basis. The incubation was carried out with rumen inoculum from two lactating Swedish Red cows cannulated in the rumen, fed ad libitum on a diet of 600 g/kg grass silage and 400 g/kg concentrate on DM basis. Diet samples were incubated for 48 h in 60 ml of buffered rumen fluid (rumen fluid:buffer ratio of 1:4 by volume) as described by Chagas et al. (2019). The *in vitro* batch fermentation was run in a fully automated system that allows continuous recording of gas production (Ramin and Huhtanen, 2012).

Methane production, acetate, butyrate, propionate, and ammonia were measured throughout the incubation period. Methane was measured at 0, 2, 4, 8, 24, 36 and 48 h according to (Ramin and Huhtanen, 2012). Gas production was measured using a fully automated system (Gas Production Recorder, GPR-2, Version 1.0 2015, Wageningen UR), with readings made every 12 min and corrected to the normal air pressure (101.3 kPa). Methane concentration was determined with a Varian Star 3400 CX gas chromatograph (Varian Analytical Instruments, Walnut Creek, CA, USA) equipped with a thermal conductivity detector. The volatile fatty acids (VFAs) were measured at 0, 8, 24 and 48 h and determined using a Waters Alliance 2795 UPLC system as described by (Puhakka *et al*., 2016). Ammonia was measured at 0 and 24h and analysed with a continuous flow analyzer (AutoAnalyzer 3 HR, SEAL Analytical Ltd., Southampton, UK) and according to the method provided by SEAL Analytical (Method no. G-102-93 multitest MT7). For model calibration, we only considered data until 24h, since microbial fermentation stopped around this time.

### 2.2. Mathematical modelling

We adapted the mathematical model of *in vitro* rumen fermentation developed by (Muñoz-Tamayo *et al*., 2016) to account for the effect of *A. taxiformis* on the fermentation. This model represents the rumen microbiota by three microbial functional groups (sugar utilisers, amino acid utilisers and methanogens). Hexose monomers are represented by glucose and amino acids are represented by an average amino acid. The model is an aggregated representation of the anaerobic digestion process that comprises the hydrolysis of cell wall carbohydrates (NDF - Neutral Detergent Fiber), non-fiber carbohydrates (NSC – Non Structural Carbohydrates) and proteins, the fermentation of soluble monomers producing the VFAs acetate, butyrate, propionate, and the hydrogenotrophic methanonogenesis. The original model was calibrated using *in vitro* experimental data from (Serment et al., 2016). Figure 1 displays a schematic representation of the rumen fermentation model indicating the effect of *A. taxiformis* on the fermentation. We assumed that microbial cells are formed by proteins and non-fiber carbohydrates and that dead microbial cells are recycled as carbon sources in the fermentation.

**Figure 1.**
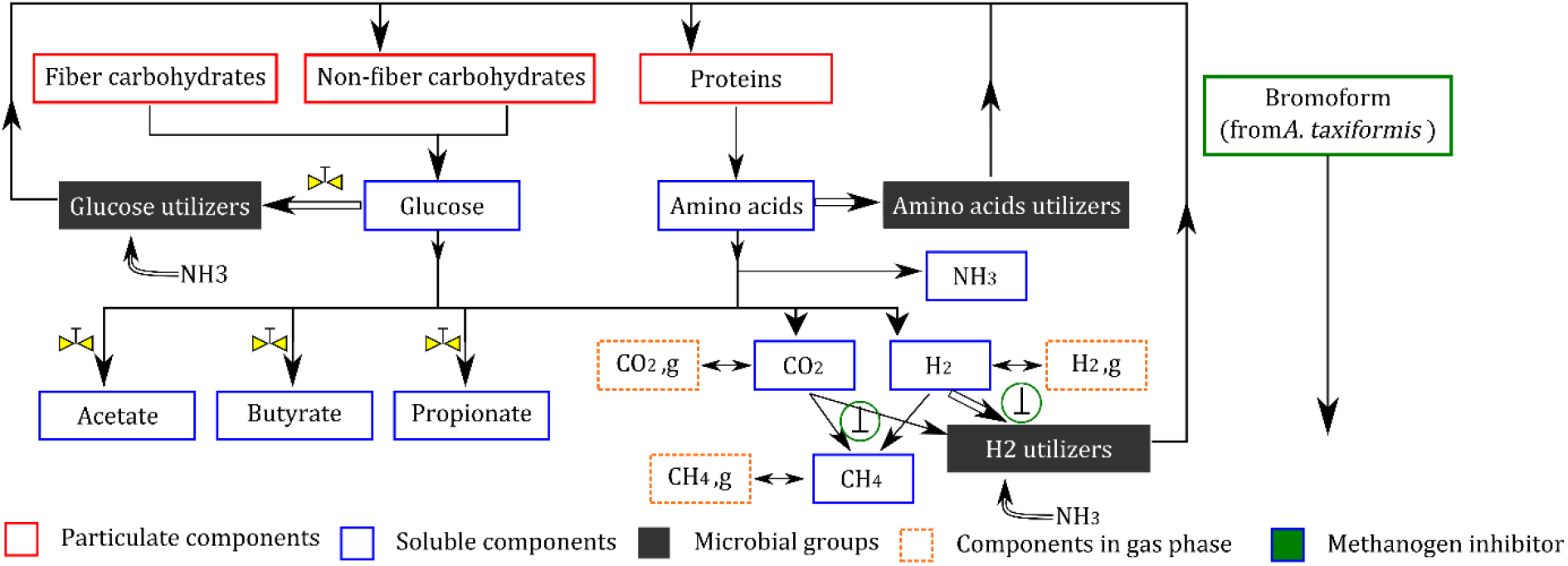
Representation of the rumen fermentation model (adapted from (Muñoz-Tamayo *et al*., 2016)). Hydrolysis of carbohydrates (fiber and non-fiber) and proteins releases respectively sugars and amino acids soluble monomers which are further utilized by the microbiota. The utilization of substrate is directed to product formation (single arrows) and microbial growth (double arrows). Each substrate is utilized by a single microbial functional group. The bromoform contained in *A. taxiformis* produces a direct inhibition of the growth rate of methanogens that results in a reduction of methane production and in an accumulation of hydrogen. The symbol 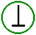 indicates the direct effect of the bromoform on the methanogenesis. Hydrogen exerts control on sugars utilization and on the flux allocation towards volatile fatty acids production. The symbol 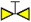 indicates the hydrogen control effect on the rumen fermentation.

The model is derived from mass balance equations of a closed system under the assumption that the protocol of gas sampling does not affect substantially the dynamics of methane and fermentation dynamics. Our model is described in compact way as follows

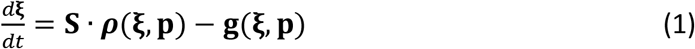

Where ξ is the vector of state variables (metabolites), ***ρ***(·) is a vector function with the kinetic rates of hydrolysis and substrate (sugars, amino acids, hydrogen) utilization. Hydrolysis rates are described by first-order kinetics. Substrate utilization rates are described by the Monod kinetics. **S** is the stoichiometry matrix containing the yield factors (*Y_i,j_*) of each metabolite (*i*) for each reaction (*j*), **g**(·) is a vector function with the equations representing transport phenomena (liquid–gas transfer), and **p** is the vector of the model parameters. The original model has 18 state variables (compartments in Figure. 1) and was implemented in Matlab (the code is accessible at https://doi.org/10.5281/zenodo.4047640). An implementation in R software is also available (Kettle *et al*., 2018). In the present work, we incorporated an additional state variable to represent the dynamics of bromoform concentration. The original model was extended to account for the impact of *A. taxiformis* on the rumen fermentation. While the original model predicts the pH, we set the pH value to 6.6.

The impact of *A. taxiformis* on the fermentation and methane production was ascribed to two mechanisms, namely the (i) direct inhibition of the growth rate of methanogens by bromoform and (ii) hydrogen control on sugars utilization and on the flux allocation towards volatile fatty acids production. These aspects are detailed below.

For the methanogenesis, the reaction rate of hydrogen utilization *ρ*_H_2__ (mol/(L h)) is given by

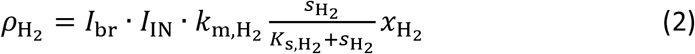

where *S*_H_2__ (mol/L) is the hydrogen concentration in liquid phase, *x*_H_2__, (mol/L) is the concentration of hydrogen-utilizing microbes (methanogens), *k*_m,H_2__ (mol/(mol h)) is the maximum specific utilization rate constant of hydrogen and *K*_s,H_2__ (mol/L) is the Monod affinity constant of hydrogen utilization, and *I*_IN_ is a nitrogen limitation factor. The kinetic rate is inhibited by the anti-methanogenic compounds of *A. taxiformis*. The factor *I*_br_ represents this inhibition as function of the bromoform concentration. We used the following sigmoid function to describe *I*_br_

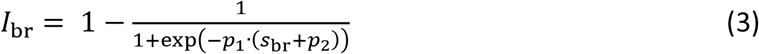

where *S*_br_ is the bromoform concentration (g/L) and *p*_1_, *p*_2_ are the parameters of the sigmoid function. We included in our model the dynamics of bromoform using a first-order kinetics to take into account that the inhibition of *A. taxiformis* declines on time as a result of the consumption of anti-methanogenic compounds (Kinley *et al*., 2016). The dynamics of *S*_br_ is

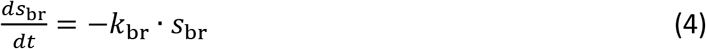

where *k*_br_ (1/h) is the kinetic rate constant of bromoform utilization.

With regard to sugars utilization, we assumed that the effect of *A. taxiformis* is ascribed to hydrogen control due to accumulation of hydrogen resulting from the methanogenesis inhibition. Hydrogen level influences the fermentation pattern (Janssen, 2010). We used the structure proposed by (Mosey, 1983) to account for hydrogen control on sugar utilization and flux allocation. However, we used different parametric functions to those proposed by (Mosey, 1983). The functions proposed by (Mosey, 1983) did not provide satisfactory results. In our model, the kinetic rate of sugar utilization is described by

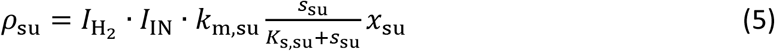

where *S*_su_ (mol/L) is the concentration of sugars, *x*_m,su_ (mol/L), is the concentration of sugar utilisers microbes, *K*_s,su_ (mol/(mol h) is the maximum specific utilization rate constant of sugars and *K*_s,su_ (mol/L) is the Monod affinity constant of sugars utilization.

The factor *I*_H_2__ describes the hydrogen inhibition:

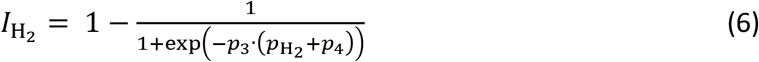

with *p*_H_2__ the hydrogen partial pressure (*p*_H_2__).

In our model, the rumen fermentation is represented by the macroscopic reactions in Table 1.

**Table 1.** Macroscopic reactions used in our model to representing rumen fermentation. For the anabolic reactions of microbial formation, we assume that microbial biomass has the molecular formula C_5_H_7_O_2_N.

|  |  |
| --- | --- |
| *Sugars (glucose) utilization* |  |
| $C_6H_{12}O_6 + 2H_2O \rightarrow 2CH_3COOH + 2CO_2 + 4H_2$ | R<sub>1</sub> |
| $3C_6H_{12}O_6 \rightarrow 2CH_3COOH + 4CH_3CH_2COOH + 2CO_2 + 2H_2O$ | R<sub>2</sub> |
| $C_6H_{12}O_6 \rightarrow CH_3CH_2CH_2COOH + 2CO_2 + 2H_2$ | R<sub>3</sub> |
| $5C_6H_{12}O_6 + 6NH_3 \rightarrow 6C_5H_7O_2N + 18H_2O$ | R<sub>4</sub> |
| *Amino acid utilization* |  |
| $C_5H_9.8O_{2.7}N_2 \rightarrow Y_{IN,aa}NH_3 + (1 - Y_{aa}) \cdot \sigma_{ac,aa} CH_3COOH + (1 - Y_{aa}) \cdot \sigma_{pr,aa} CH_3CH_2COOH + (1 - Y_{aa}) \cdot \sigma_{bu,aa} CH_3CH_2CH_2COOH + (1 - Y_{aa}) \cdot \sigma_{IC,aa} CO_2 + (1 - Y_{aa}) \cdot \sigma_{H_2,aa} H_2 + Y_{aa}C_5H_7O_2N$ | R<sub>5</sub>* |
| *Hydrogen utilization* |  |
| $4H_2 + 2CO_2 \rightarrow CH_4 + 2H_2O$ | R<sub>6</sub> |
| $10H_2 + 5CO_2 + NH_3 \rightarrow C_5H_7O_2N + 8H_2O$ | R<sub>7</sub> |

\*R<sub>5</sub> is an overall reaction resulting from weighing the fermentation reactions of individual amino acids.

Table 1 shows that VFA production from glucose utilization occurs *via* reactions R_1_-R_3_. The pattern of the fermentation is determined by the flux allocation of glucose utilization through these three reactions. We denote *λ_k_* as the molar fraction of the sugars utilized *via* reaction *k*. It follows that *λ*_1_ + *λ*_2_ + *λ*_3_ = 1.

The fermentation pattern (represented in our model by the flux allocation parameters *λ*_k_) is controlled by thermodynamic conditions and by electron-mediating cofactors such as nicotinamide adenine dinucleotide (NAD) that drive anaerobic metabolism via the transfer of electrons in metabolic redox reactions (Mosey, 1983; Hoelzle *et al*., 2014; van Lingen *et al*., 2019). In our model, the regulation exerted by the NADH/NAD+ couple on the flux allocation is incorporated *via* regulation functions that are dependent on the hydrogen partial pressure (*p*_H_2__). This hybrid approach resulted by assuming a linearity between the couple NADH/NAD+ and the *p*_H_2__ following the work of (Mosey, 1983; Costello *et al*., 1991). As discussed by (van Lingen *et al*., 2019), the production of acetate *via* the reaction R_1_ is favoured at low NADH/NAD+ while the production of propionate *via* the reaction R_2_ is favoured at high NADH/NAD+. Accordingly, we represented the flux allocation parameters by the following sigmoid functions:

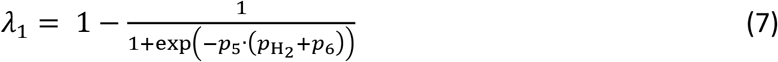

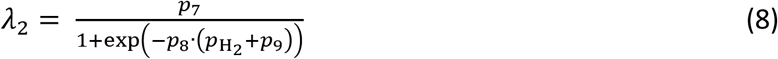

Our model then predicts that high levels of supplementation of *A. taxiformis* will result in high hydrogen levels that will favour propionate production (R_2_) over acetate production (R_1_). By this parameterization of the flux allocation parameters, our model accounts for the concomitant reduction of the acetate:propionate ratio that is observed when methane production is reduced.

### 2.3. Parameter estimation

We used the maximum likelihood estimator that minimizes the following objective function

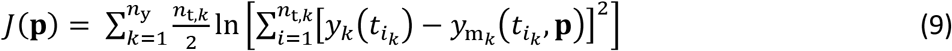

Where **p** is the vector of parameters to be estimated, *n_y_* is the number of measured variables, *n*_t,*k*_ is the number of observation times of the variable *k*, *t*_i_*k*__ is the *i*th measurement time for the variable *y_k_*, and *y*_m_k__ is the value predicted by the model. The measured variables are the concentrations of acetate, butyrate, propionate, NH_3_, and the moles of methane produced.

We used the IDEAS Matlab^®^ toolbox (Muñoz-Tamayo *et al*., 2009) (freely available at http://genome.jouy.inra.fr/logiciels/IDEAS) to generate the function files for solving the optimization problem locally. Then, we used the generated files by IDEAS to look for global optimal solutions using the Matlab optimization toolbox MEIGO (Egea *et al*., 2014) that implements the enhanced scatter search method developed by (Egea *et al*., 2010) for global optimization.

We reduced substantially the number of parameters to be estimated by setting most of the model parameters to the values reported in the original model implementation and using the information obtained from the *in vitro* study (Chagas *et al*., 2019). For example, the hydrolysis rate constant for NDF was obtained from (Chagas *et al*., 2019) whereas the hydrolysis rate constants of NSC (*k*_hydr,nsc_) and proteins (*k*_hydr,pro_) were included in the parameter estimation problem. The kinetic rate constant for hydrogen utilization *k*_m,H_2__ was set 16 mol/(mol h) using an average value of the values we obtained for the predominant archaea *Methanobrevibacter ruminantium* and *Methanobrevibacter smithii* (Muñoz-Tamayo *et al*., 2019) using a microbial yield factor of 0.006 mol biomass/mol H_2_ (Pavlostathis *et al*., 1990). With this strategy, we penalize the goodness-of-fit of the model. But, on the other hand, we reduce practical identifiability problems typically found when calibrating biological kinetic-based models (Vanrolleghem *et al*., 1995). The parameter vector for the estimation is then **p**: {*k*_hydr,nsc_, *k*_hydr,pro_, *k*_br_, *p*_1_,*p*_2_,…*p*_9_}. The optimization was set in a multi-experiment fitting context that integrates the data of all treatments. To evaluate the model performance, we computed the determination coefficient (R^2^), the Lin’s concordance correlation coefficient (CCC) (Lin, 1989), the Root mean squared error (RMSE) and the coefficient of variation of the RMSE (CV_RMSE_). We also performed residual analysis for bias assessment according to (St-Pierre, 2003).

## 3. Results

### 3.1. Dynamic prediction of rumen fermentation

The extended model developed in the present work to account for the impact of *A. taxiformis* on the rumen fermentation is freely available at https://doi.org/10.5281/zenodo.4090332 with all the detailed information of the model and the experimental data used for model calibration. An open source version in the Scilab software (https://www.scilab.org/) was made available to facilitate reproductibility since Scilab files can be opened with a text editor. The software R (https://www.r-project.org/) was used to plot the figures.

Figure 2 shows the dynamic data of fermentation variables for the levels of *A. taxiformis* at 0.06% and 0.5% compared against the model predicted variables. Figure 3 displays the comparison of all observations against model predictions. Figure 4 shows the residuals for all variables against centred predicted values.

**Figure 2.**
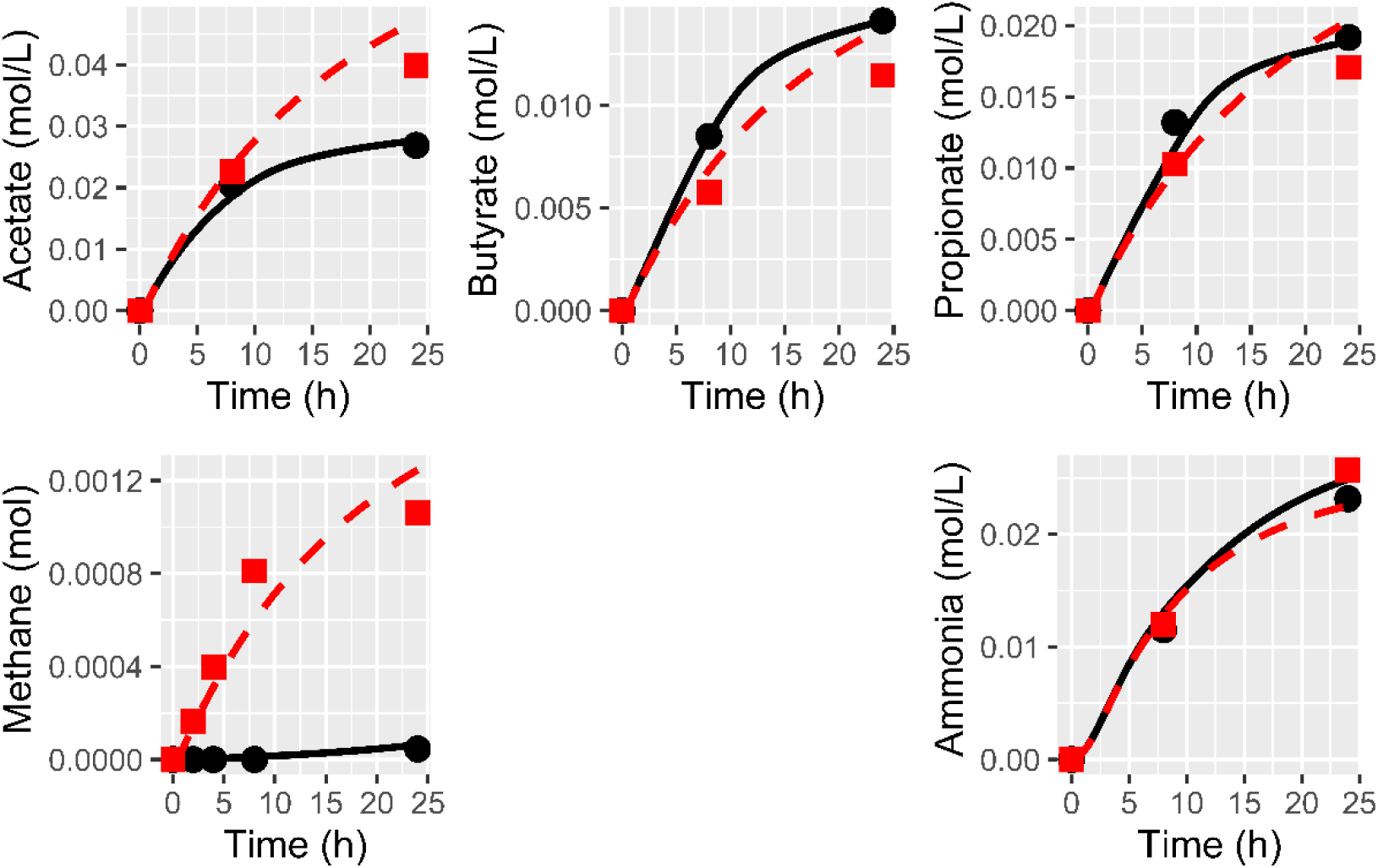
Example of model fitting. Experimental data of fermentation variables for the levels of *A. taxiformis* at 0.5% (•) and 0.06% 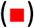 are compared against the model predicted responses in solid black lines (for 0.5% level) and in dashed red lines (for the 0.06% level).

**Figure 3.**
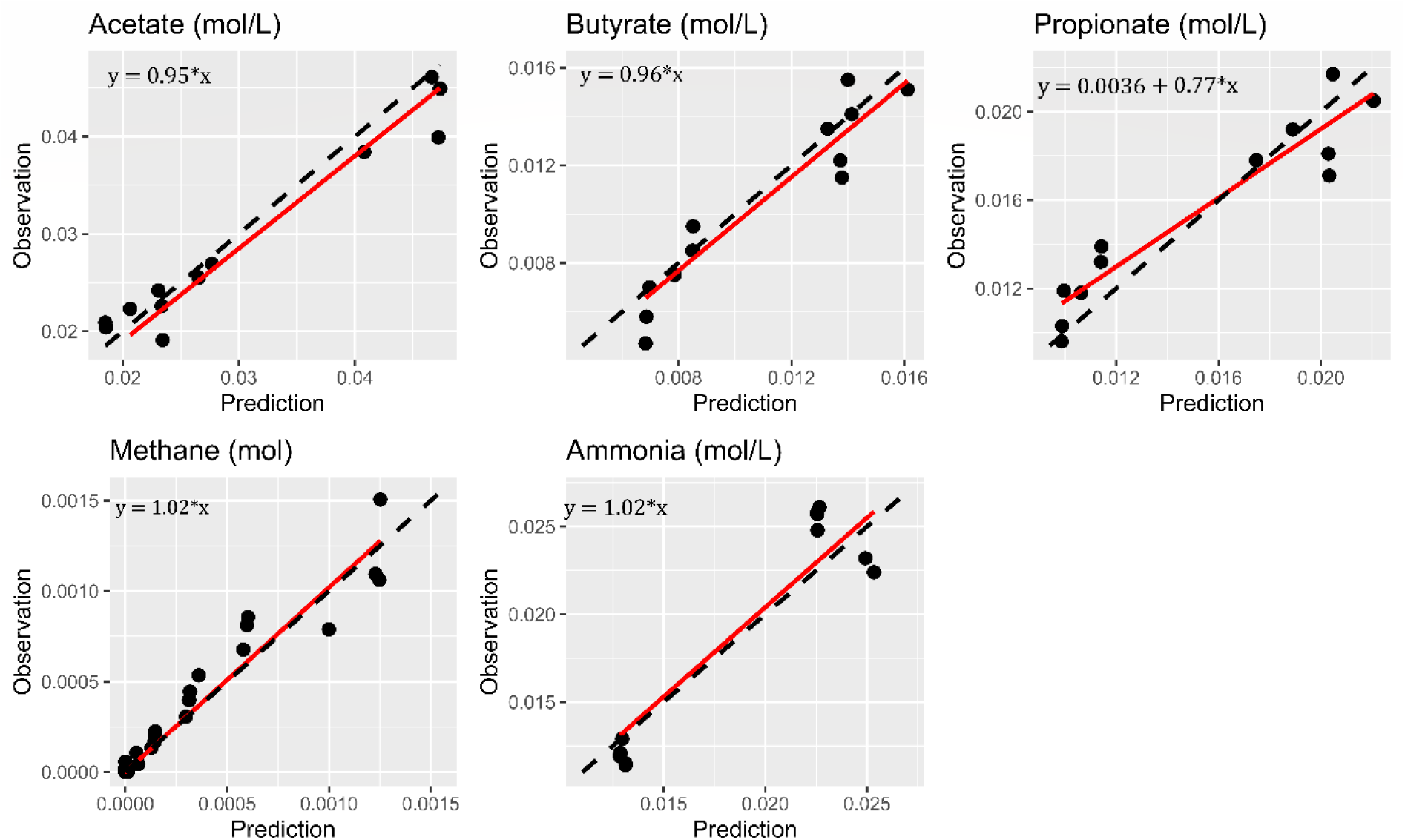
Summary of the model performance calibration integrating data of all treatments. Experimental data (•) are plotted against the model predicted variables. Solid lines are the linear fitted curve. Dashed lines are the isoclines. Only the intercept of the curve of propionate was different from zero at 5% significance level.

**Figure 4.**
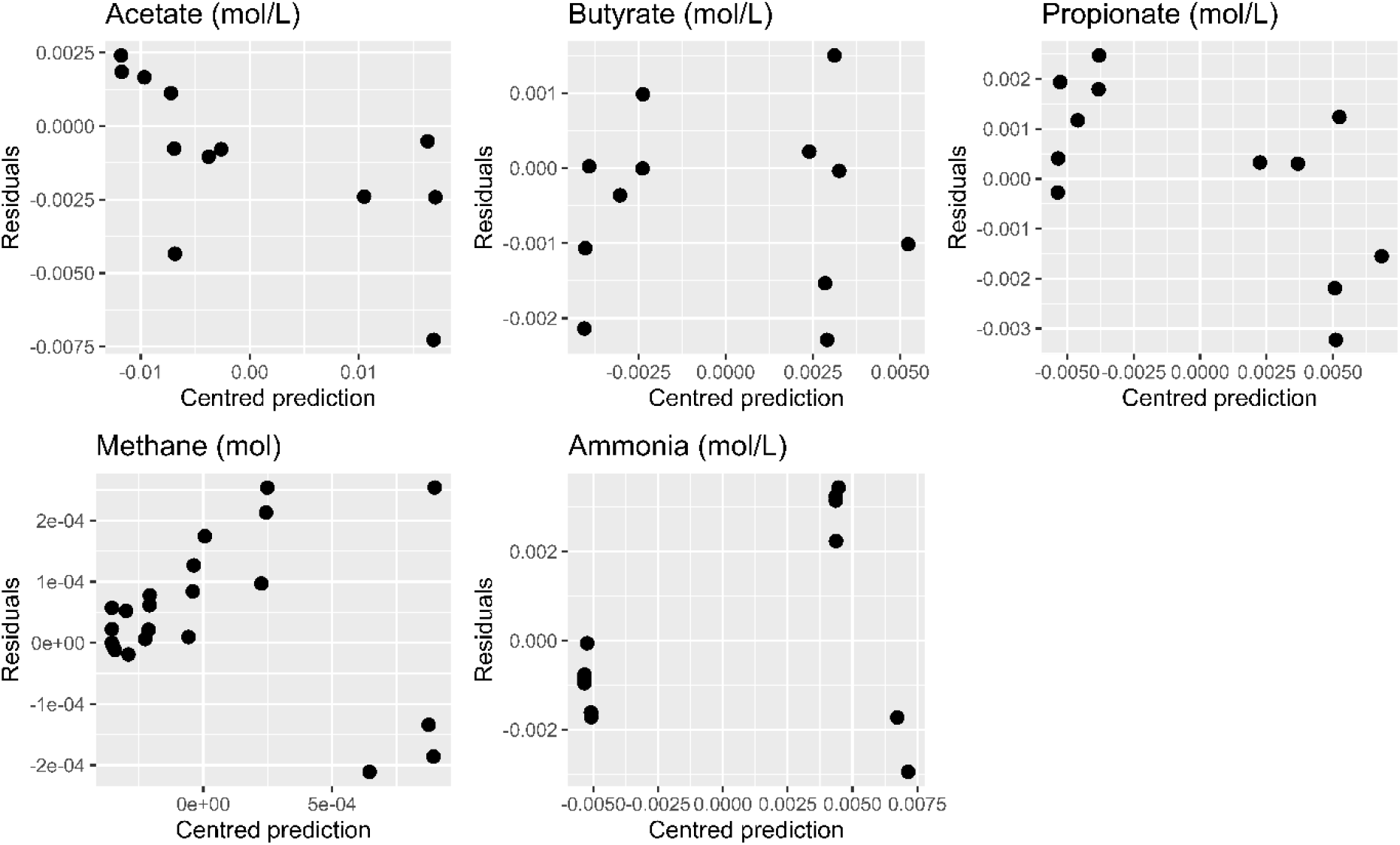
Residuals values of observed variables against centred predicted variables (*n*_CH4_ =24, *n*_NH3_ = *n*_ac_ = *n*_bu_ = *n*_pr_ = 12).

To evaluate the performance of our model and its validation, external independent data is required. Due to data limitation, we did not perform such a validation. To provide indicators of our model, we calculated standard statistical indicators of model performance which are shown in Table 2. These statistic indicators are biased and thus should be looked with caution since they are calculated using the calibration data. Nevertheless, they provide an indication of the adequacy of the model structure to represent the fermentation dynamics. For methane, butyrate and NH3 the mean and linear biases were not significant at the 5% significance level. Acetate and propionate exhibited significant linear bias. The liquid compounds have an average coefficient of variation of the RMSE (CV(RMSE)) of 11.25%. Methane had the higher CV(RMSE) (31%). The concordance correlation coefficients were higher than 0.93. Propionate had the lowest determination coefficient (R^2^=0.82) while methane and the other compounds had a R^2^ close to 0.9.

**Table 2.** Statistical indicators of model performance.

|  | Acetate | Butyrate | Propionate | Methane | NH<sub>3</sub> |
| --- | --- | --- | --- | --- | --- |
| R<sup>2</sup> | 0.91 | 0.88 | 0.82 | 0.92 | 0.89 |
| RMSE<sup>a</sup> | 0.0029 | 0.0012 | 0.0017 | 1.21x10<sup>-4</sup> | 0.002 |
| 100×CV<sub>RMSE</sub><sup>b</sup> | 10 | 12 | 11 | 31 | 12 |
| CCC<sup>c</sup> | 0.96 | 0.94 | 0.93 | 0.96 | 0.93 |
| Residual analysis |  |  |  |  |  |
| $residual = \alpha + \beta \cdot (predicted - mean\ predicted\ value)$ | | | | | |
|  | Acetate | Butyrate | Propionate | Methane | NH<sub>3</sub> |
| $\alpha$ (p-value) | -0.0010<br>(p= 0.14) | -0.00047<br>(p= 0.21) | 0.00019<br>(p= 0.63) | 4.0e-05<br>(p= 0.12) | 0.00012<br>(p= 0.86) |
| $\beta$ (p-value) | -0.15<br>(p= 0.024) | 0.0028<br>(p= 0.98) | -0.22<br>(p= 0.024) | -0.031<br>(p= 0.60) | 0.15<br>(p= 0.23) |

<sup>a</sup> Root mean squared error (RMSE).<sup>b</sup> Coefficient of variation of the RMSE (CV(RMSE)).<sup>c</sup> Concordance correlation coefficient (CCC)

Figure 5 compares specifically the experimental data of methane against the model predictions for all levels of *A. taxiformis*.

**Figure 5.**
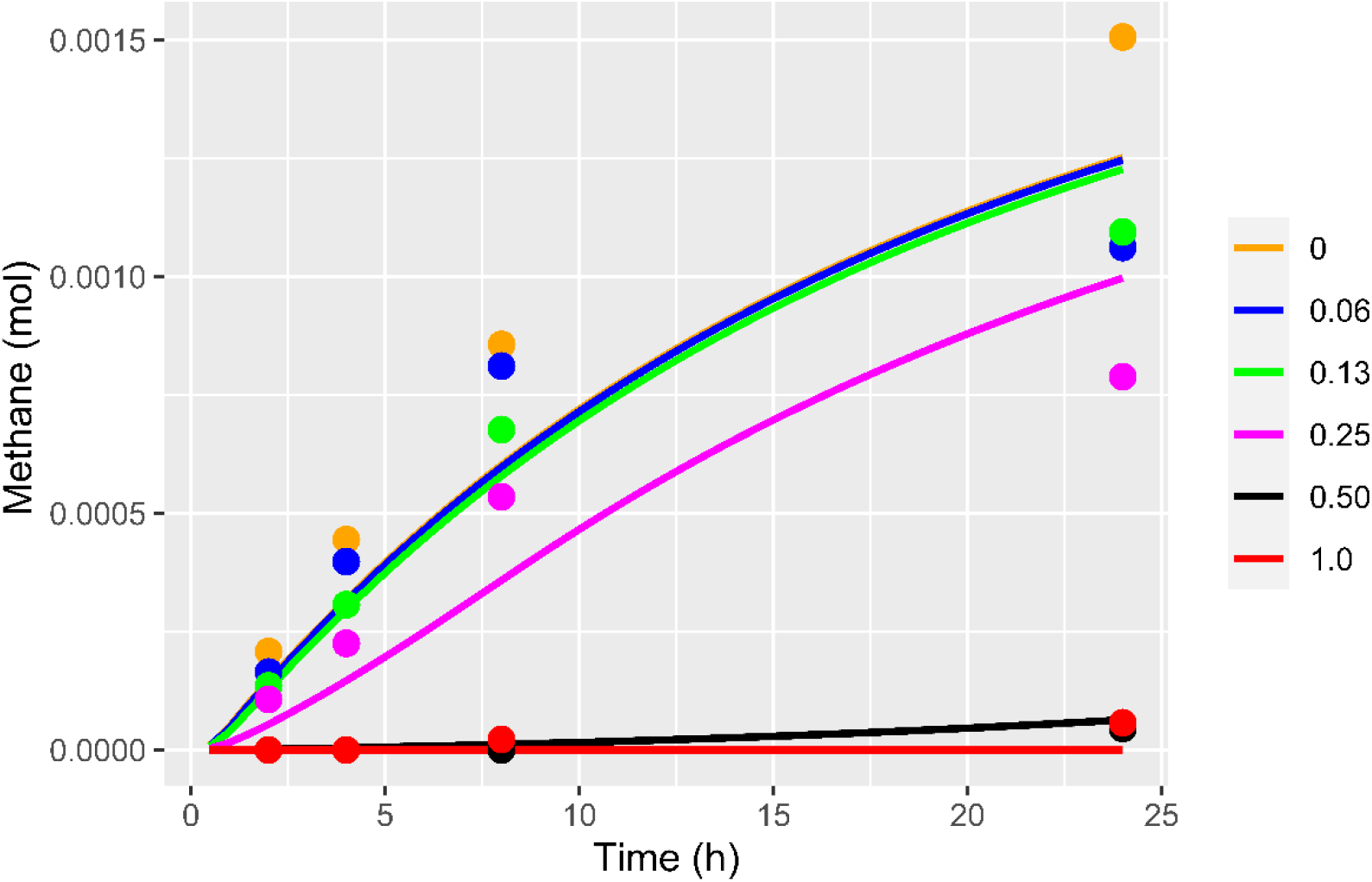
Experimental observations of methane (circles) in the headspace of the incubation system are compared against the model predicted values (solid lines). Increase of the dose of *A. taxiformis* results in a decrease of methane production.

### 3.2. Prediction of the factors representing the impact of *A. taxiformis* on rumen fermentation

Figure 6 plots the factors that represent the effect of *A. taxiformis* on rumen fermentation. Direct inhibition of the methanogenesis due to the anti-methanogenic action of bromoform is represented by the factor *I*_br_. Methanogenesis inhibition results in hydrogen accumulation impacting the flux allocation of sugars utilization.

**Figure 6.**
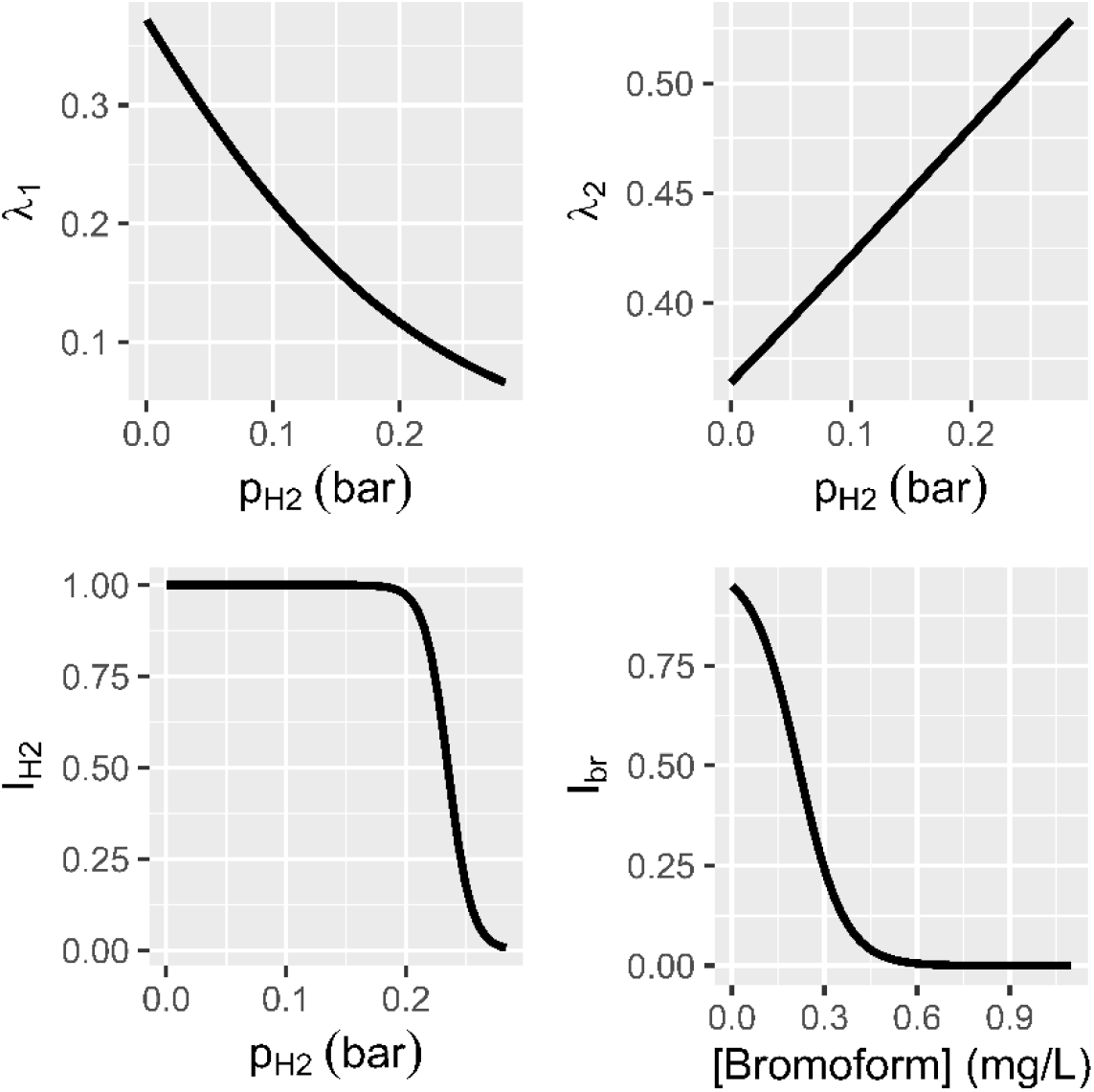
In our model, the effect of *A. taxiformis* on rumen fermentation is represented by a direct inhibitory effect of bromoform (*I*_br_) on the methanogens growth rate. Methanogenesis inhibition results in hydrogen accumulation. Hydrogen control impacts sugar utilization by inhibiting the rate of sugar utilization (factor *I*_H_2__) and by regulating the flux allocation parameters (*λ*_1_, *λ*_2_) towards VFA production.

Figure 7 displays the simulated dynamics of hydrogen in the headspace for all the supplementation levels of *A. taxiformis*. For supplementation levels higher than 0.25%, the methanogenesis inhibition resulted in a substantial hydrogen accumulation.

**Figure 7.**
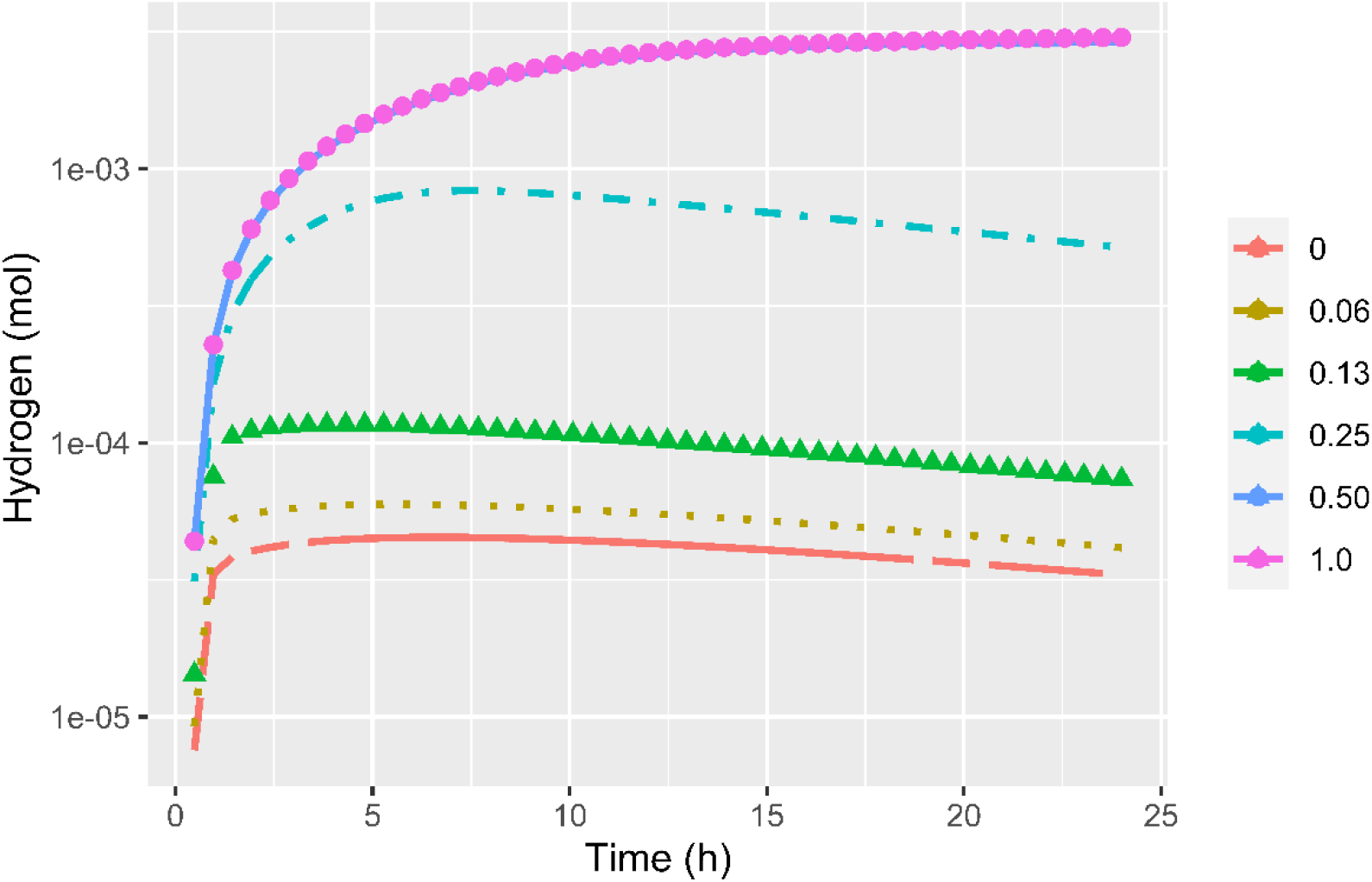
Predicted dynamics of hydrogen in the headspace for levels of *A. taxiformis*. Increase of the dose of *A. taxiformis* results in an increase of hydrogen in the incubation system.

## 4. Discussion

The goal of this work was to model the impact of *A. taxiformis* supplementation on the rumen microbial fermentation and methane production under *in vitro* conditions using experimental data from (Chagas *et al*., 2019). Overall, our model was able to capture the dynamics of VFA, ammonia and methane production for different levels of *A. taxiformis* indicating the potential of the model structure towards the development of predictive models for assessing methane mitigation strategies in ruminants. With the exception of propionate, the slope of observed vs predicted variables is very close to one. Model limitations will be discussed further. We modelled the effect of *A. taxiformis* on rumen fermentation by two mechanisms. The first mechanism is associated to the direct inhibition of the methanogens growth rate by the anti-methanogenic compounds of *A. taxiformis* documented in different studies (Kinley *et al*., 2016; Machado *et al*., 2016a; Roque *et al*., 2019). In our model, we ascribed the inhibitory effect of *A. taxiformis* only to the concentration of bromoform. The first-order kinetic rate for bromoform consumption and the inhibition factor (*I*_br_) (Fig. 6) allowed our model to account for the observed dynamic decline in methanogenesis inhibition (Kinley *et al*., 2016). It should be noted that although bromoform is the most abundant anti-methanogenic compound in *A. taxiformis*, the anti-methanogenic capacity of *A. taxiformis* is the result of the synergetic action of all halogenated products present in the macroalgae (Machado *et al*., 2016b). Accordingly, it will be useful to include further in our model other secondary compounds such as dibromochloromethane. To enhance our model, it will be central to perform new experiments to characterize the dynamics of anti-methanogenic compounds. This aspect is of great relevance to allow the model to be adapted to different applications of seaweed supplementation since it is known that the composition of halogenic compounds can vary with respect to the season, harvesting and drying methods.

The second mechanism that accounts for the impact of *A. taxiformis* on the fermentation is hydrogen control, which it is discussed below.

### 4.1. Methane inhibition and hydrogen control

The anti-methanogenic capacity of *A. taxiformis* is dose-dependent. The experimental study of (Chagas *et al*., 2019) showed that methane production was inhibited almost completely by *A. taxiformis* at a level of 0.5%. Our model predictions aligned with the experimental observations (Fig. 5). The anti-methanogenic capacity of the macroalgae leads to hydrogen accumulation (Kinley *et al*., 2020; Roque *et al*., 2020) as predicted by our model in Fig. 7. The level of hydrogen increases as the dose of *A. taxiformis* increases. The predicted values of hydrogen levels in the headspace for low doses of *A. taxiformis* showing in Figure 7 are in agreement with *in vitro* reported values (Serment et al., 2016). The level of hydrogen can impact electron-mediating cofactors such as nicotinamide adenine dinucleotide (NAD) which are important drivers of anaerobic metabolism *via* the transfer of electrons in metabolic redox reactions (Hoelzle et al., 2014). van Lingen et al., 2019 extended the rumen model developed by (Dijkstra *et al*., 1992) to incorporate the regulation of NADH/NAD+ on the fermentation.

In our model, the regulation of NADH/NAD+ was incorporated *via* the control of hydrogen partial pressure assuming a linearity between the couple NADH/NAD+ and the ***p*_H_2__** and following the model structure proposed by (Mosey, 1983) with a different parameterisation for the functions describing the effect of ***p*_H_2__** on the rate of glucose utilization and on the flux allocation. The linearity assumption between NADH/NAD+ and the ***p*_H_2__** might not be fulfilled for all values of ***p*_H_2__**(De Kok et al., 2013). In the experimental conditions used in the experiment here analysed (Chagas et al., 2019) and under rumen physiological conditions, the linearity between NADH/NAD+ might be valid.

With regard to the hydrogen control on glucose utilization, our model predicts that the inhibition is effective at ***p*_H_2__** higher than 0.2 bar (factor *I*_H_2__ in Fig. 6). In our model, the incorporation of the inhibitory effect of hydrogen was motivated to account for the decrease of the total production of VFA at high levels of supplementation of *A. taxiformis* observed by (Chagas *et al*., 2019). Such a decrease of VFA production is dose-dependent as observed in *in vitro* studies (Kinley *et al*., 2016; Machado *et al*., 2016a). *In vivo*, while insignificant changes in total VFA concentration between a control diet and diets with *A. taxiformis* supplementation were observed in Brangus steers (Kinley *et al*., 2020), inclusions of *A. taxiformis* resulted in a decrease in total VFA ruminal concentration in sheep compared with control diet (Li *et al*., 2016). Accordingly, additional studies with simultaneous measurements of VFA and hydrogen are needed to validate the relevance of the inhibitory term *I*_H_2__ of our model both under *in vitro* and *in vivo* conditions.

In addition to the impact of *A. taxiformis* supplementation on methane reduction, it is important to look at the effects on animal productivity. *A. taxiformis* impacts the production of VFAs, which are energy sources for the animal. Accordingly, changes in VFA production might result in changes on productivity and feed efficiency. Optimal feeding strategies should thus be designed to attain a trade-off between low methane emissions and high productivity and animal health. Studies showing the effect of *A. taxiformis* supplementation on live weight (Li *et al*., 2016), average daily weight gain and feed conversion efficiency (Kinley *et al*., 2020; Roque *et al*., 2020) are still scarce to provide a large data base for concluding on the impact of *A. taxiformis* on animal productivity and feed efficiency. However, the studies of (Kinley *et al*., 2020; Roque *et al*., 2020) suggest that feed conversion efficiency tend to increase concomitantly with the reduction of methane production induced by an adequate supplementation of *A. taxiformis*, supporting the theory of redirection of energy otherwise lost as methane (Kinley *et al*., 2020). An opportunity to enhance the action of *A. taxiformis* might be the implementation of a feeding strategy integrating macroalgae supplementation with an adequate additive allowing to redirect metabolic hydrogen towards nutritional fermentation products beneficial to the animal. Such a strategy will fulfil the objectives of reducing methane emissions while increasing animal productivity (Ungerfeld, 2020).

With regard to the fermentation pattern, when the hydrogen level increases the hydrogen control operates by increasing the flux of carbon towards propionate (*λ*_2_) while the flux towards the reaction that produces only acetate (*λ*_1_) decreases (Fig. 6). Incorporating hydrogen control on the fermentation pattern in our model enabled us to predict the decrease of the acetate to propionate ratio observed at levels of *A. taxiformis* supplementation leading to substantial methane reduction both *in vitro* (Machado *et al*., 2016a; Chagas *et al*., 2019) and *in vivo* (Kinley *et al*., 2020). Our model is also consistent with *in vitro* (Kinley *et al*., 2016; Machado *et al*., 2016a) and *in vivo* (Stefenoni *et al*., 2021) studies showing the increase of butyrate level when the inclusion of *A. taxiformis* increases.

### 4.2. Model limitations and perspectives

In our model, the quantification of the impact of *A. taxiformis* was ascribed by the action of bromoform on the methanogens growth rate and by the action of *p*_H_2__ on the fermentation pattern. However, in the experimental study of (Chagas *et al*., 2019), nor bromoform nor *p*_H_2__ were measured. From our bibliography search, we did not find studies reporting dynamic measurements of bromoform. Although we did not perform an identifiably analysis, we might expect that the lack of bromoform and hydrogen data in our work might result in structural identifiability (Muñoz-Tamayo *et al*., 2018) and model distinguishability problems (Walter and Pronzato, 1996). We will then require external data to validate our model. Experiments to be done within the MASTER project (https://www.master-h2020.eu/contact.html) will fill this gap and provide data for challenging and improving our model.

Our model aligns with the efforts of enhancing the dynamic prediction of ruminal metabolism *via* the incorporation of thermodynamics and regulation factors (Offner and Sauvant, 2006; Ghimire *et al*., 2014; van Lingen *et al*., 2019). While our work focused only on hydrogen control on sugars metabolism, future work is needed to incorporate the impact of *A. taxiformis* supplementation on amino acids fermentation. The study of (Chagas *et al*., 2019) showed a decrease of branched-chain volatile fatty acids (BCVFA) with increased supplementation of *A. taxiformis*. Such a decrease of BCVFA might have a negative influence on microbial activity.

We modelled the regulation of sugars metabolism by hydrogen control following a grey-box modelling approach where the regulation factors were assigned to sigmoid functions without an explicit mechanistic interpretation. However, to enhance the understanding of rumen fermentation, it will be useful to pursue an approach incorporating the role of internal electron mediating cofactors on the direction of electrons towards hydrogen or VFA (Hoelzle *et al*., 2014; Ungerfeld, 2020). Recent progress in this area (van Lingen *et al*., 2019) opens a direction for improving the prediction of rumen models.

The ultimate goal of this work is to pursue a model extension to account for *in vivo* conditions. In this endeavour, experimental data in semi-continuous devices such as the Rusitec (Roque *et al*., 2019a) will be instrumental for model improvement. *In vivo*, in addition to the impact on fermentation, *A. taxiformis* can induce changes in rumen mucosa (Li *et al*., 2016). These mucosa changes might translate in changes on the rate of absorption of ruminal VFA. This effect on the rate of VFA absorption should be quantified and incorporated into an extended model. In our model, the pH was set constant. However, pH exhibits a dynamic behaviour that can impact the activity of the rumen microbiota. The impact of the pH on the rumen microbial groups should be then considered in a future version of the model, integrating the mechanistic calculation of pH elaborated in our previous model (Muñoz-Tamayo *et al*., 2016).

Finally, although our model developments focused on the impact of *A. taxiformis* on rumen fermentation and methane production, we think our model structure has the potential to be applied to other additives such as 3-nitrooxypropanol (Hristov *et al*., 2015; Duin *et al*., 2016) whose action is specifically directed to inhibit methanogenic archaea, as the halogenated compounds of *A. taxiformis*. We expect these model developments can be useful to help the design of sustainable nutritional strategies promoting healthy rumen function and low environmental footprint.

## 5. Conclusions

We have developed a rumen fermentation model that accounts for the impact of *A. taxiformis* supply on *in vitro* rumen fermentation and methane production. Our model was effective in representing the dynamics of VFA, ammonia and methane for six supplementation levels of *A. taxiformis*, providing a promising prediction tool for assessing the impact of additives such as seaweeds on rumen microbial fermentation and methane production in *vitro*.

## 6. Declarations

### Ethics approval

Not applicable

### Availability of data and material

The datasets and codes used in this study are available at https://doi.org/10.5281/zenodo.4090332

## Acknowledgements

Authors thank Henk van Lingen (Wageningen University, The Netherlands), Alberto Atzori (University of Sassari, Italy) and two anonymous reviewers appointed by Luis Tedeschi (Texas A&M University, USA) for the evaluation of this manuscript by PCI Animal Science (https://animsci.peercommunityin.org/). Their reviews have greatly improved this paper. Version 4 of this preprint has been peer-reviewed and recommended by Peer Community In Animal Science (https://doi.org/10.24072/pci.animsci.100006).

## Authors’ contributions

JCC, MH and SJC produced the experimental data of the study. RMT developed the mathematical model and drafted the article. All authors contributed to the analysis and interpretation of the results. All authors read and approved the final manuscript.

## References

Beauchemin, K.A., Ungerfeld, E.M., Eckard, R.J., and Wang, M. (2020) Review: Fifty years of research on rumen methanogenesis: Lessons learned and future challenges for mitigation. Animal 14(S1): S2–S16.

Chagas, J.C., Ramin, M., and Krizsan, S.J. (2019) In vitro evaluation of different dietary methane mitigation strategies. Animals 9: 1120.

Chalupa, W. (1977) Manipulating Rumen Fermentation. J Anim Sci 46: 585–599.

Costello, D.J., Greenfield, P.F., and Lee, P.L. (1991) Dynamic Modeling of a Single-Stage High-Rate Anaerobic Reactor. 1. Model Derivation. Water Res 25: 847–858.

Czerkawski, J.W. and Breckenridge, G. (1975) New inhibitors of methane production by rumen micro-organisms. Development and testing of inhibitors in vitro. Br J Nutr 34: 429–446.

Denman, S.E., Tomkins, N.W., and McSweeney, C.S. (2007) Quantitation and diversity analysis of ruminal methanogenic populations in response to the antimethanogenic compound bromochloromethane. FEMS Microbiol Ecol 62: 313–322.

Dijkstra, J., Neal, H.D., Beever, D.E., and France, J. (1992) Simulation of nutrient digestion, absorption and outflow in the rumen: model description. J Nutr 122: 2239–2256.

Dubois, B., Tomkins, N.W., D. Kinley, R., Bai, M., Seymour, S., A. Paul, N., and Nys, R. de (2013) Effect of Tropical Algae as Additives on Rumen in Vitro Gas Production and Fermentation Characteristics. Am J Plant Sci 4: 34–43.

Duin, E.C., Wagner, T., Shima, S., Prakash, D., Cronin, B., Yáñez-Ruiz, D.R., et al. (2016) Mode of action uncovered for the specific reduction of methane emissions from ruminants by the small molecule 3-nitrooxypropanol. Proc Natl Acad Sci U S A 113: 6172–6177.

Egea, J.A., Henriques, D., Cokelaer, T., Villaverde, A.F., MacNamara, A., Danciu, D.P., et al. (2014) MEIGO: An open-source software suite based on metaheuristics for global optimization in systems biology and bioinformatics. BMC Bioinformatics 15: 136.

Egea, J.A., Martí, R., and Banga, J.R. (2010) An evolutionary method for complex-process optimization. Comput Oper Res 37: 315–324.

Ellis, J.L., Dijkstra, J., France, J., Parsons, A.J., Edwards, G.R., Rasmussen, S., et al. (2012) Effect of high-sugar grasses on methane emissions simulated using a dynamic model. J Dairy Sci 95: 272–285.

Evans, F.D. and Critchley, A.T. (2014) Seaweeds for animal production use. J Appl Phycol 26: 891–899.

Ghimire, S., Gregorini, P., and Hanigan, M.D. (2014) Evaluation of predictions of volatile fatty acid production rates by the Molly cow model. J Dairy Sci 97: 354–362.

Hoelzle, R.D., Virdis, B., and Batstone, D.J. (2014) Regulation mechanisms in mixed and pure culture microbial fermentation. Biotechnol Bioeng 111: 2139–2154.

Hristov, A.N., Oh, J., Giallongo, F., Frederick, T.W., Harper, M.T., Weeks, H.L., et al. (2015) An inhibitor persistently decreased enteric methane emission from dairy cows with no negative effect on milk production. Proc Natl Acad Sci U S A 112: 10663–10668.

Huws, S.A., Creevey, C.J., Oyama, L.B., Mizrahi, I., Denman, S.E., Popova, M., et al. (2018) Addressing global ruminant agricultural challenges through understanding the rumen microbiome: past, present, and future. Front Microbiol 9: 2161.

Janssen, P.H. (2010) Influence of hydrogen on rumen methane formation and fermentation balances through microbial growth kinetics and fermentation thermodynamics. Anim Feed Sci Technol 160: 1–22.

Kettle, H., Holtrop, G., Louis, P., and Flint, H.J. (2018) microPop: Modelling microbial populations and communities in R. Methods Ecol Evol 9: 399–409.

Kinley, R.D., Martinez-Fernandez, G., Matthews, M.K., de Nys, R., Magnusson, M., and Tomkins, N.W. (2020) Mitigating the carbon footprint and improving productivity of ruminant livestock agriculture using a red seaweed. J Clean Prod 259: 120836.

Kinley, R.D., De Nys, R., Vucko, M.J., MacHado, L., and Tomkins, N.W. (2016) The red macroalgae Asparagopsis taxiformis is a potent natural antimethanogenic that reduces methane production during in vitro fermentation with rumen fluid. Anim Prod Sci 56: 282–289.

Li, X., Norman, H.C., Kinley, R.D., Laurence, M., Wilmot, M., Bender, H., et al. (2016) Asparagopsis taxiformis decreases enteric methane production from sheep. Anim Prod Sci 58: 681–688.

Lin, L.I. (1989) A concordance correlation-coefficient to evaluate reproducibility. Biometrics 45: 255–268.

van Lingen, H.J., Fadel, J.G., Moraes, L.E., Bannink, A., and Dijkstra, J. (2019) Bayesian mechanistic modeling of thermodynamically controlled volatile fatty acid, hydrogen and methane production in the bovine rumen. J Theor Biol 480: 150–165.

Machado, L., Magnusson, M., Paul, N.A., Kinley, R., de Nys, R., and Tomkins, N. (2016a) Dose-response effects of Asparagopsis taxiformis and Oedogonium sp. on in vitro fermentation and methane production. J Appl Phycol 28: 1443–1452.

Machado, L., Magnusson, M., Paul, N.A., Kinley, R., de Nys, R., and Tomkins, N. (2016b) Identification of bioactives from the red seaweed Asparagopsis taxiformis that promote antimethanogenic activity in vitro. J Appl Phycol 28: 3117–3126.

Machado, L., Magnusson, M., Paul, N.A., De Nys, R., and Tomkins, N. (2014) Effects of marine and freshwater macroalgae on in vitro total gas and methane production. PLoS One 9: e85289.

Machado, L., Tomkins, N., Magnusson, M., Midgley, D.J., de Nys, R., and Rosewarne, C.P. (2018) In Vitro Response of Rumen Microbiota to the Antimethanogenic Red Macroalga Asparagopsis taxiformis. Microb Ecol 75: 811–818.

Maia, M.R.G., Fonseca, A.J.M., Oliveira, H.M., Mendonça, C., and Cabrita, A.R.J. (2016) The potential role of seaweeds in the natural manipulation of rumen fermentation and methane production. Sci Rep 6: 32321.

Makkar, H.P.S., Tran, G., Heuzé, V., Giger-Reverdin, S., Lessire, M., Lebas, F., and Ankers, P. (2016) Seaweeds for livestock diets: A review. Anim Feed Sci Technol 212: 1–17.

Mosey, F.E. (1983) Mathematical-Modeling of the Anaerobic-Digestion Process - Regulatory Mechanisms for the Formation of Short-Chain Volatile Acids from Glucose. Water Sci Technol 15: 209–232.

Muñoz-Tamayo, R., Giger-Reverdin, S., and Sauvant, D. (2016) Mechanistic modelling of in vitro fermentation and methane production by rumen microbiota. Anim Feed Sci Technol 220: 1–21.

Muñoz-Tamayo, R., Laroche, B., Leclerc, M., and Walter, E. (2009) IDEAS: A parameter identification toolbox with symbolic analysis of uncertainty and its application to biological modelling. In, IFAC Proceedings Volumes., pp. 1271–1276.

Muñoz-Tamayo, R., Popova, M., Tillier, M., Morgavi, D.P., Morel, J.P., Fonty, G., and Morel-Desrosiers, N. (2019) Hydrogenotrophic methanogens of the mammalian gut: Functionally similar, thermodynamically different—A modelling approach. PLoS One 14: e0226243.

Muñoz-Tamayo, R., Puillet, L., Daniel, J.B., Sauvant, D., Martin, O., Taghipoor, M., and Blavy, P. (2018) Review: To be or not to be an identifiable model. Is this a relevant question in animal science modelling? Animal 12: 701–712.

Offner, A. and Sauvant, D. (2006) Thermodynamic modeling of ruminal fermentations. 55: 343–365.

Paul, N.A., De Nys, R., and Steinberg, P.D. (2006) Chemical defence against bacteria in the red alga Asparagopsis armata: Linking structure with function. Mar Ecol Prog Ser 306: 87–101.

Pavlostathis, S.G., Miller, T.L., and Wolin, M.J. (1990) Cellulose Fermentation by Continuous Cultures of Ruminococcus-Albus and Methanobrevibacter-Smithii. Appl Microbiol Biotechnol 33: 109–116.

Puhakka, L., Jaakkola, S., Simpura, I., Kokkonen, T., and Vanhatalo, A. (2016) Effects of replacing rapeseed meal with fava bean at 2 concentrate crude protein levels on feed intake, nutrient digestion, and milk production in cows fed grass silage–based diets. J Dairy Sci 99: 7993–8006.

Ramin, M. and Huhtanen, P. (2012) Development of an in vitro method for determination of methane production kinetics using a fully automated in vitro gas system-A modelling approach. Anim Feed Sci Technol 174: 190–200.

Roque, B.M., Brooke, C.G., Ladau, J., Polley, T., Marsh, L.J., Najafi, N., et al. (2019) Effect of the macroalgae Asparagopsis taxiformis on methane production and rumen microbiome assemblage. Anim Microbiome 1:3:

Roque, B.M., Salwen, J.K., Kinley, R., and Kebreab, E. (2019) Inclusion of Asparagopsis armata in lactating dairy cows’ diet reduces enteric methane emission by over 50 percent. J Clean Prod 234: 132–138.

Roque, B.M., Venegas, M., Kinley, R., DeNys, R., Neoh, T.L., Duarte, T.L., et al. (2020) Red seaweed (Asparagopsis taxiformis) supplementation reduces enteric methane by over 80 percent in beef steers. bioRxiv.

St-Pierre, N.R. (2003) Reassessment of Biases in Predicted Nitrogen Flows to the Duodenum by NRC 2001. J Dairy Sci 86: 344–350.

Stefenoni, H.A., Räisänen, S.E., Cueva, S.F., Wasson, D.E., Lage, C.F.A., Melgar, A., et al. (2021) Effects of the macroalga Asparagopsis taxiformis and oregano leaves on methane emission, rumen fermentation, and lactational performance of dairy cows. J Dairy Sci.

Ungerfeld, E.M. (2020) Metabolic Hydrogen Flows in Rumen Fermentation: Principles and Possibilities of Interventions. Front Microbiol 11: 589.

Vanrolleghem, P.A., Vandaele, M., and Dochain, D. (1995) Practical identifiability of a biokinetic model of activated-sludge respiration. Water Res 29: 2561–2570.

Walter, E. and Pronzato, L. (1996) On the identifiability and distinguishability of nonlinear parametric models. Math Comput Simul 42: 125–134.

Wang, Y., Xu, Z., Bach, S.J., and McAllister, T.A. (2008) Effects of phlorotannins from Ascophyllum nodosum (brown seaweed) on in vitro ruminal digestion of mixed forage or barley grain. Anim Feed Sci Technol 145: 375–395.

Wood, J.M., Kennedy, F.S., and Wolfe, R.S. (1968) The Reaction of Multihalogenated Hydrocarbons with Free and Bound Reduced Vitamin B12. Biochemistry 7: 1707–1713.

